# Open ocean and coastal strains of the N_2_-fixing cyanobacterium UCYN-A have distinct transcriptomes

**DOI:** 10.1101/2022.07.26.501530

**Authors:** María del Carmen Muñoz-Marín, Jonathan D. Magasin, Jonathan P. Zehr

**Affiliations:** Ocean Sciences Department, University of California, Santa Cruz, California, United States of America; Departamento de Bioquímica y Biología Molecular, Campus de Excelencia Internacional Agroalimentario, Universidad de Córdoba, Córdoba, Spain

## Abstract

Decades of research on marine N_2_ fixation focused on *Trichodesmium*, which are generally free-living cyanobacteria, but in recent years the endosymbiotic cyanobacterium *Candidatus* Atelocyanobacterium thalassa (UCYN-A) has received increasing attention. However, few studies have shed light on the influence of the host versus the habitat on UCYN-A N_2_ fixation and overall metabolism. Here we compared transcriptomes from natural populations of UCYN-A from oligotrophic open-ocean versus nutrient-rich coastal waters, using a microarray that targets the full genomes of UCYN-A1 and UCYN-A2 and known genes for UCYN-A3. We found that UCYN-A2, usually regarded as adapted to coastal environments, was transcriptionally very active in the open ocean and appeared to be less impacted by habitat change than UCYN-A1. Across habitats and sublineages, genes for N_2_ fixation and energy production had high transcript levels, and, intriguingly, were among the minority of genes that kept the same schedule of diel expression. This might indicate different regulatory mechanisms for genes that are critical to the symbiosis for the exchange of nitrogen for carbon from the host. Our results underscore the importance of N_2_ fixation in UCYN-A symbioses across habitats, with consequences for community interactions and global biogeochemical cycles.

## Introduction

Ocean productivity depends on the availability of nitrogen (N). Disolved atmospheric N_2_, though plentiful in surface waters, is biologically unavailable to the marine microorganisms that drive ocean productivity. Instead, they largely rely on fixed inorganic N compounds such as ammonium and nitrate, which they rapidly deplete to low levels in the well-lit surface waters of the open ocean gyres. Only some microorganisms can fix N_2_, reducing it to ammonium, and are the main source of N for productivity in terrestrial and aquatic environments (1). In the marine environment several species of cyanobacteria, including symbionts of diatoms and haptophytes, are known to be important N_2_-fixers. These include the uncultivated unicellular cyanobacteria *Candidatus* Atelocyanobacterium thalassa (UCYN-A) because of their high N_2_ fixation and growth rates (2–4) and potential for N transfer into the food web through grazers (2, 5, 6). DNA sequencing showed that UCYN-A has a massively streamlined genome (7, 8) that lacks the machinery required for carbon fixation, including photosystem II [PSII] and RuBisCO, as well as other key metabolic pathways, e.g. the entire tricarboxylic acid (TCA) cycle (7, 9, 10). Later it was discovered that UCYN-A lives in symbiosis with a photosynthetic haptophyte, related to *Braarudosphaera bigelowii*, with which it exchanges fixed N for fixed carbon (9, 10). Subsequent work showed that disrupting photosynthesis by the host arrested UCYN-A cell division and changed its daily pattern of *nifH* transcription (11). This indicates a tight coupling between UCYN-A and the host, but the interplay of environmental and host-mediated factors on the metabolism and distribution of UCYN-A are not understood.

Phylogenetic analyses of one of the nitrogenase genes (*nifH*), the key enzyme in N_2_ fixation, have shown that there are at least five distinct UCYN-A sublineages (12–15). The symbioses are globally distributed and include nutrient-enriched coastal waters (3, 16–21), oligotrophic open ocean (2, 15, 22, 23), and low-temperature waters (24–26). Some UCYN-A sublineages occur more frequently than others on sinking particles (27), which might impact particle-associated communities and the export of fixed N_2_. All of this suggests that UCYN-A diversity and distribution are important for a complete picture of N_2_ fixation in the global ocean.

It has been shown that UCYN-A fixes N_2_ during the day and coordinates N_2_ fixation and general metabolism similarly to other cyanobacteria that fix N_2_ during the day such as *Trichodesmium* and heterocyst-forming cyanobacteria (16). Critical to this coordination is the protection of nitrogenase from oxygen evolved by photosynthesis by the host, because oxygen irreversibly damages nitrogenase (28, 29). Although much gene expression in cyanobacteria follows circadian rhythms that are driven by three “clock” (*kai*) genes (30–32), UCYN-A lacks two of the three *kai* genes. Thus, UCYN-A’s mechanism for coordinating expression with that of its host is poorly understood (33). UCYN-A might be controlled by a novel circadian network or the circadian rhythm of the host, and/or UCYN-A might simply respond to host metabolites (16).

Many essential aspects of the symbioses have not yet been explored, in particular, the metabolism under different nutrient conditions. If UCYN-A depends on metabolite production by the host, does its metabolism differ in nutrient-rich versus poor waters? Metatranscriptomes for UCYN-A1 and UCYN-A2 from nutrient-rich coastal waters were previously analyzed (16). The present work examines transcriptional patterns in oligotrophic water (characterized by low concentrations of macronutrients and warm surface temperatures). Specifically, we analyzed transcription over diel cycles by UCYN-A sublineages A1, A2, and A3 at an open-ocean station in the North Pacific Subtropical Gyre (NPSG) and, for UCYN-A1 and A2, compared to diel transcriptional patterns in coastal waters. In the open ocean, for all three sublineages ~11% of detected genes had cyclic expression patterns with maxima and minima levels that mainly occurred at or shortly after sunrise or sunset. In contrast, in coastal waters ~85% of detected genes from UCYN-A1 and UCYN-A2 were cyclically expressed with maxima or minima that seldom occurred near sunrise or sunset. Different nutrient and light conditions compared to the coastal study might have influenced the different expression patterns. However, at both sites many of the same genes were highly transcribed, in particular genes that support N_2_ fixation. Our findings offer clues to UCYN-A1/2/3 habitat preferences and regulation by their respective hosts in open-ocean versus coastal environments, just when culture studies for some UCYN-A sublineages have become possible.

## Materials and Methods

### *In situ* diel sampling of UCYN-A

Samples were collected from CTD casts at Stn. ALOHA (22° 45’ N, 158° 0’ W at 45 m) during the C-MORE Cruise C-20 (http://hahana.soest.hawaii.edu/hot/cruises.html) between 6th and 10th April, 2015. Seawater was collected in 12-L polyvinylchloride bottles affixed to a rosette sampler equipped with Sea-Bird 911+ conductivity, temperature, and pressure sensors.

A total of 16 samples covered a full light-dark cycle with 8 time points and 3 h intervals. Time points were as follows: 9:00-L3, 12:00-L6, 15:00-L9, 18:00-L12, 21:00-D3, 00:00-D6, 03:00-D9, 06:00-2D12, 09:00-2L3 and 12:00-2L6, where L and D indicate the light and dark period, respectively, 2L and 2D the second light-dark cycle, and the number the corresponding hours into the light or dark period. Two replicates were collected for each time point except for D3, D6, as well as L3 and 2L3.

Samples were collected by filtering a total of 500 mL from each seawater replicate through 0.22 μm pore-size, 47 mm diameter Supor filters (Pall Corporation, Port Washington, NY, USA) using a peristaltic pump. Filters were placed in sterile 2 mL bead-beating tubes with sterile glass beads, flash-frozen in liquid nitrogen and stored at −80°C until extraction.

### RNA extraction and processing for hybridization to the microarray

Environmental RNA containing transcripts from UCYN-A cells was extracted using the Ambion RiboPure Bacteria Kit (Ambion^®^, ThermoFisher) and treated Turbo-DNA-free^™^ DNase Kit (Ambion^®^, ThermoFisher) following the procedure described in (16).

RNA purity, concentration and quality were determined using a NanoDrop 1000 (Thermo Scientific, Waltham, MA, USA) and a 2100 Bioanalyzer (Agilent Technologies, Santa Clara, CA, USA) using the RNA 6000 Nano kit (Agilent Technologies). Only samples with RNA Integrity Number >7.0 and ratios of A260/A230 and A260/A280 ≥1.8 were processed further.

Double-stranded (ds) cDNA was synthesized and amplified following the procedure described in Muñoz-Marín et al., 2019. The amplified cDNA was purified with the GenElute PCR cleanup kit (Sigma-Aldrich), and the quality and quantity of ds-cDNA was determined with NanoDrop 1000 and a 2100 Bioanalyzer using the Agilent DNA 7500 kit (Agilent Technologies). Four hundred ng of total RNA yielded on average 12 μg of ds-cDNA. The labeling and hybridization of cDNA samples (1.0 μg of ds-cDNA) to the microarray was done at the Roy J. Carver Center for Genomics (CCG) Facility (University of Iowa, Iowa city, Iowa, USA) according to the Agilent Technology protocol for arrays.

### Design of the UCYN-A array

The UCYN-A oligonucleotide expression array targeted all known genes from UCYN-A1 and UCYN-A2 with oligonucleotide probe sequences (“probes”) from the array design described in (16). For UCYN-A3 314 genes were targeted, all of them reconstructed in (15). The UCYN-A3 probes were designed with the eArray web-based tool (Agilent Technology Inc.; https://earray.chem.agilent.com/earray/) using a similar approach to that described in (16) for UCYN-A1 and UCYN-A2.

Briefly, 1602 probes, each 60 nucleotides (nt) in length, were designed for the 314 UCYN-A3 genes. Each gene was usually represented by 6 distinct probes. The UCYN-A1, A2, and A3 probes were tested *in silico* for possible cross-hybridization to non-UCYN-A taxa. Candidate probes were rejected if they had BLASTN alignments that were ≥ 95% nt identity over the full probe length to non-UCYN-A sequences. These alignment criteria simulated the hybridization specificity of Agilent SurePrint technology. Non-UCYN-A sequences were from nucleotide databases from the Community Cyberinfrastructure for Advanced Microbial Ecology Research and Analysis (CAMERA (34), now available in iMicrobe (35)) in June 2014 (Marine microbes, Microbial Eukaryote Transcription and Non-redundant Nucleotides).

Next, candidate probes that passed the cross-hybridization filter were checked for UCYN-A sublineage specificity. Probes were clustered using CD-HIT-EST (36, 37) at 95% nt similarity. A probe was rejected if it clustered with probes from other UCYN-A sublineages. A few exceptions were made for probes from highly conserved genes (such as the nitrogenase gene, *nifH*).

In summary, 6088 probes for 1195 UCYN-A1genes, 6324 probes for 1244 UCYN-A2 genes, and 1351 probes for 314 UCYN-A3 genes were chosen. Each of these 13,763 total experimental probes were replicated 4 times at different microarray coordinates (55,052 experimental features). There were also 1319 standard control probes defined in Agilent Technology Array (IS-62976-8-V2_60Kby8_GX_EQC_201000210) and ERCC control probes) included at 7924 random feature locations on the microarray. The final design of the microarray included ca. 62976 features and was synthesiszed as 8 arrays per slide, two slides for our 16 total samples.

The probe sequences are available at NCBI Gene Expression Omnibus (GEO) under accession number GSE206403.

### Microarray analyses

Microarray data were prepared using the `MicroTOOLs` R package (https://www.jzehrlab.com/microtools) which has been used previously to analyze marine microbial community metatranscriptomes targeted by the MicroTOOLs array design (38, 39). In the present study we substituted in the UCYN-A microarray design (NCBI GEO accession GPL32341). All 16 microarrays passed quality checks from the `arrayQualityMetrics` R package v3.44 (40). Probe intensities were normalized across samples by quantiles (41) and then converted to gene intensities (“transcript levels” in Results, log_2_ scale) using robust multi-array averaging (42) using the `affyPLM` R package v1.64 (43). Then genes were detected within each sample if they had a signal-to-noise ratio (SNR, or *z*-score) that was >5, with noise defined as the median intensity of the bottom 10% of genes in the sample. A total of 2753 genes were detected based on SNR>5. However, we also required detected genes to be higher than the least concentrated spike-in control (ERCC-00048) in >3 samples. This additional requirement was satisfied based on either of two comparisons: (i) array intensities of the gene versus the ERCC; (ii) predicted transcript counts of the gene versus the ERCC, based on a linear model. After filtering based on ERCC spike-ins, the total number of detected genes was 1939 (S1 Table).

To identify genes with significantly periodic transcript levels (“diel genes”; false discovery rate <0.25), we used the `fdrfourier` function in the R package `cycle` v1.42 with AR1 background models (44) and 1000 background data sets. 188 diel genes were identified at Stn. ALOHA. After standardizing transcript levels (0 mean, unit variance), diel genes were clustered with the `pvclust` R package v2.2 (correlation based distances, clustering by centroids, 1000 bootstraps, cluster significance alpha = 0.95; (45)). Assigned clusters are indicated in S1 Table, with most diel genes in either a cluster with sunrise peaks (D12, cluster "sunrise_peak_edge.172") or sunset peaks (L12, cluster “sunset_peak_edge.174”).

Additional tests indicated that the number of diel genes was contingent on habitat, and that fewer diel genes were identified at Stn. ALOHA even after controlling for different numbers of time points and detected genes in comparison to the coastal study (Supplementary Information).

Previously published metatranscriptomic data from South Atlantic TARA Oceans Stn. 78 was used, specifically the metagenome-normalized sequencing read counts in Tables S1, S2 of Cornejo-Castillo et al. (2016). Gene loci were used to map genes in the TARA study to GIDs in the UCYN-A microarray design, which allowed transferring pathway and other annotations from the microarray to the TARA data. To combine the Stn. 78 MiSeq data with our array data in Fig 1, quantiles were assigned to detected pathways within each sample. Pathways that were detected in more than one third of samples were retained and hierarchically clustered using `pvclust` as described above except average linkage clustering was used. Samples were hierarchically clustered the same way, but standardizing was unnecessary for quantiles.

**Fig 1.**
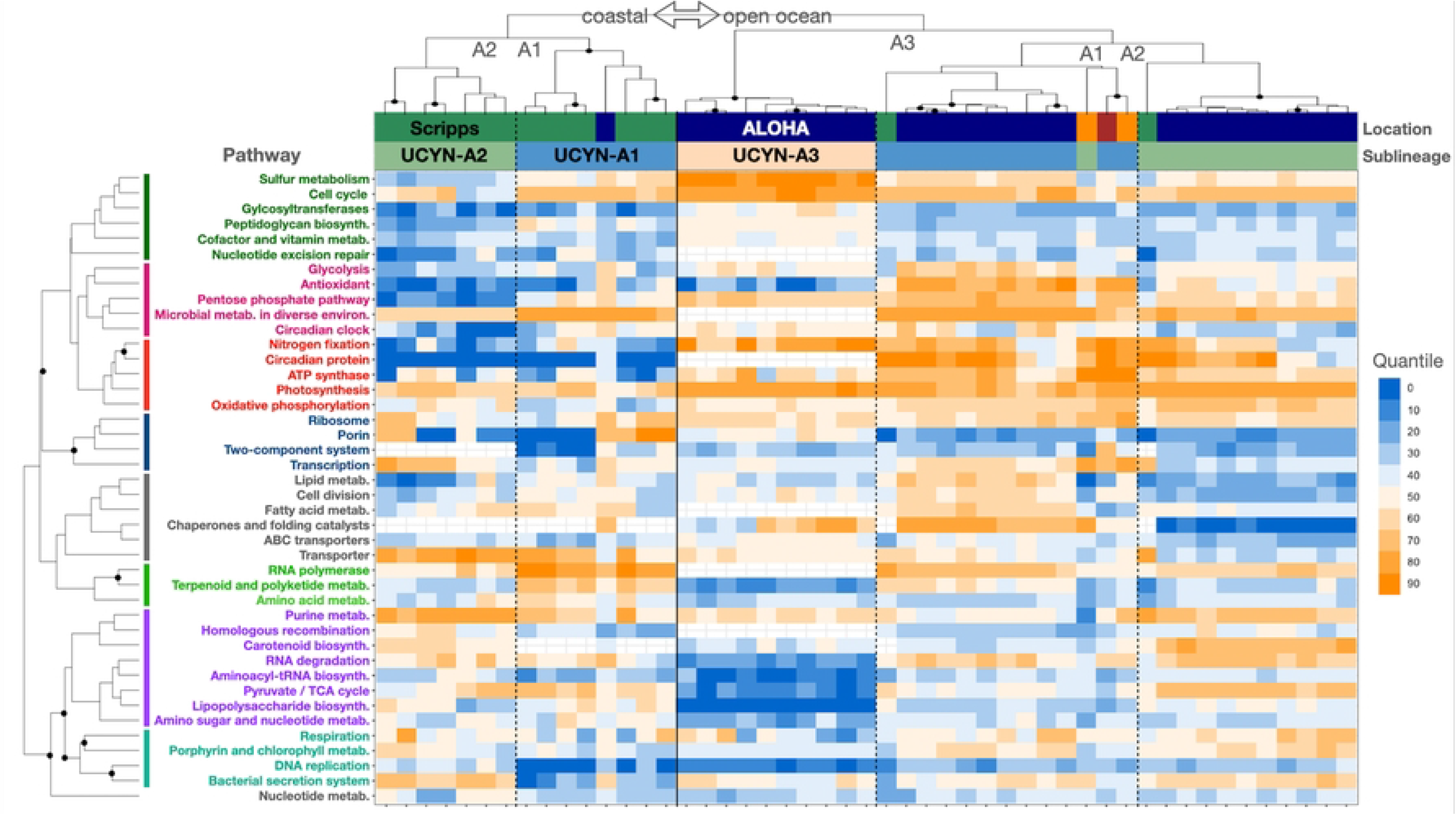
The heat map shows that expression is driven first by location (Stn. ALOHA, Scripps Pier, Tara SAtl), then by sublineage. Each column is a specific sublineage at a location and is standardized (quantiles). Cell colors are the quantile for the median of the genes in the pathway. Sample and pathway clusters with solid discs were strongly supported (Methods). In the Location bar, samples from TARA Stns. 76 and 78 in the South Atlantic are in orange.

## Results and Discussion

### UCYN-A2 had higher transcripts per cell than other sublineages at Stn. ALOHA

The UCYN-A population at Stn. ALOHA was comprised mainly of the open ocean sublineages UCYN-A1 (87% of amplified *nifH* gene sequences) and UCYN-A3 (9.4%), and very little of the coastal sublineage UCYN-A2 (~0.7%) based on a previous study that used the same samples (15). It was therefore surprising that UCYN-A2 had microarray probe intensities as high as UCYN-A1 and A3 (Fig S1) and was as robustly detected (75%, 66%, and 73% of gene targets detected for UCYN-A1, A2, and A3, respectively; S1 Table). These results are not likely explained by cross-hybridization because *in silico* simulations indicated that only ~3% of the UCYN-A2 probes could hybridize to transcripts from UCYN-A1 or UCYN-A3 using Agilent SurePrint technology (only transcripts ≥95%id over ≥95% of the probe length will hybridize; Supplementary Information). However, it is possible that the array detected a sublineage with mainly ≥95% nucleic acid identity to the Scripps Pier UCYN-A2 isolate (SIO64986) that was used to design the probes. Genetic differences between SIO64986 and the sublineage at Stn. ALOHA might explain the slightly higher frequency of low-intensity probes for UCYN-A2 compared to UCYN-A1 and A3 (Fig S2) and slightly lower percentage of gene targets detected.

The high transcript levels for UCYN-A2 despite its rarity suggest that UCYN-A2 produced more transcripts per cell than other sublineages at Stn. ALOHA. This was corroborated by surface water metatranscriptomes collected from the south Atlantic (Tara Oceans Station 78) where 1.6× more *nifH* transcript sequences came from UCYN-A2 than UCYN-A1 (in the >0.8 μm size fraction; (46). Additionally, we found that UCYN-A2 (at Stn. 78) had on average 3.8× more transcript sequences than UCYN-A1 for genes detected for both sublineages (*p*~0, one-sided paired *t*-test, *n*=272 genes). The UCYN-A2 counts were also higher for genes detected for either sublineage in any size fraction (0.2-3, >0.8, or 5-20 μm; *p*~0, one-sided unpaired *t*-test, *n_A2_*=1020, *n_A1_*=1083). Thus, in the oligotrophic NPSG and south Atlantic, UCYN-A2 cells were transcriptionally more active than UCYN-A1, which might imply higher N_2_ fixation rates.

Indeed, higher rates of N_2_ fixation by UCYN-A2 compared to UCYN-A1 have been measured in the oligotrophic tropical North Atlantic (3); the Arctic (24); and even in nutrient-rich southern California coastal waters (3, 47), where the highest UCYN-A2 N_2_ fixation rates occurred after upwelling had increased nutrient concentrations (nitrate+nitrite and phosphate) and decreased temperature (3). This suggests that higher fixation by UCYN-A2 compared to UCYN-A1 occurs across habitats and is not likely a response by the UCYN-A2 symbiosis to low nutrients. Instead, the UCYN-A2 host might require more fixed N_2_ than the UCYN-A1 host (46). Consistent with this hypothesis, the UCYN-A2 host is larger (length ~3-4 μm versus ~2 μm for the UCYN-A1 host) (48) and contains more symbionts than the UCYN-A1 host (15, 46).

### Highly transcribed genes were similar across sublineages and environments

At Stn. ALOHA, UCYN-A sublineages had highest transcript levels for the same genes and pathways including photosynthesis (*psaAB* as well as cytochrome b_6_f complex genes *petBC*), ATP synthases, N_2_ fixation (*nifDK*), ABC transporters, cell cycle (*clpP*) and division (*ftsH*), glycolysis (FBA), respiration (*coxB*), and ribosomal proteins (Figs 1 and 2; S1 Table). UCYN-A1 and A2 shared several other top genes for the metabolism of porphyrin and chlorophyll (*hemL* and an unnamed enzyme [E1.14.13.81]), oxidative phosphorylation (*ndhCK*), bacterial secretion system (*secY*), carbohydrate porins (*oprB*), RNA degradation (*dnaK*), and the circadian oscillating protein 23 (COP23). The microarray lacks these genes for UCYN-A3.

**Fig 2.**
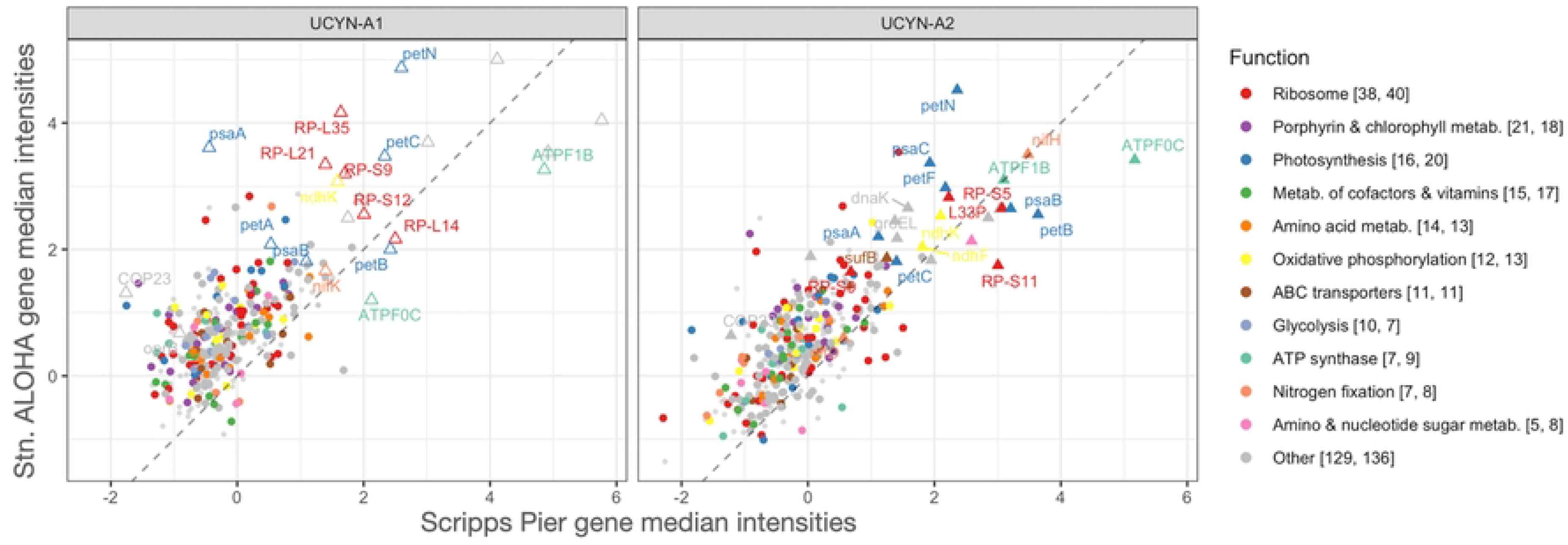
Shown are all 760 genes (*n_UCYN-A1_*=368, *n_UCYN-A2_*=392) that were detected at both Stn. ALOHA and Scripps Pier. Transcript levels were standardized (mean=0, s.d.=1) in all samples and then medians were calculated for all detected genes (*n_ALOHA_*=1939, *n_Scripps_*=762). Axes are the standardized median transcript levels and the dashed reference lines indicate equal median transcript levels at both sites. Genes are above the reference lines consistent with significantly higher median transcript levels for cross-site genes at Stn. ALOHA. Genes are colored by pathway and the legend includes in brackets the number of cross-site detected genes for UCYN-A1 and A2. Genes that had transcript levels in the top 10% at both sites for UCYN-A1 are named and shown as open triangles. Top genes for UCYN-A2 are shown with filled triangles, and names shared with UCYN-A1 indicate homologs.

The most highly transcribed genes at Stn. ALOHA and Scripps Pier were similar for UCYN-A1, as well as for UCYN-A2, and came from several of the pathways already mentioned: photosynthesis, cytochrome b_6_f complex, ATP synthases, N_2_ fixation, and oxidative phosphorylation (Fig 2). Moreover, we identified genes from these pathways among the top transcribed by both UCYN-A1 and A2 in the South Atlantic Ocean (46). The high transcript levels for genes in these pathways across sublineages and environments suggests that they are fundamental to UCYN-A symbioses, perhaps because they function in the regeneration of reductant and ATP needed for N_2_ fixation as proposed by Muñoz-Marín et al. 2019.

We also compared all detected genes at Stn. ALOHA and Scripps Pier. Far more genes were detected in the present work (1939 genes) than at Scripps Pier (762 genes; S1 Table). This is likely explained by higher UCYN-A transcript relative abundances in the oligotrophic community compared to the coastal ((49); Supplementary Information). If so, then the coastal study favored the detection of highly transcribed UCYN-A genes but missed less transcribed genes, which were detected only at Stn. ALOHA. Consistent with this hypothesis nearly every gene detected at Scripps Pier was also detected at Stn. ALOHA (760 of 762 genes), and the 760 cross-site detected genes (*n_A1_*=362 and *n_A2_*=398) had significantly higher median transcript levels than the other 1179 genes detected only at Stn. ALOHA (*p~0* in Welch’s *t*-tests performed separately for UCYN-A1 and A2; Fig 2). A similar result was seen for UCYN-A3: Transcript levels from UCYN-A3 genes that had homologs to cross-site detected genes were significantly higher (*p*~3.1E-7 in Welch’s *t*-test comparison of levels from 103 UCYN-A3 genes with cross-site homologs to 44 without). Altogether these observations indicate that (1) more of each UCYN-A metatranscriptome was detected at Stn. ALOHA than at Scripps Pier, and (2) highly transcribed genes and pathways were similar across UCYN-A sublineages and environments.

Correlation analyses also showed overall similar transcription intensities across sites but additionally suggested that the UCYN-A1 symbiosis was more sensitive to habitat (Figs 2 and S3). Each sublineage (considered separately) tended to transcribe genes in similar order of intensity across sites (Spearman’s *ρ_A1_*=0.57, *ρ_A2_*=0.70, and *p*~0 for both; Fig 2). Moreover, pathways were often highly correlated across sites (Fig S3). For example, transcript levels for ribosomal genes were highly correlated across sites for UCYN-A1 (Pearson’s *ρ*_A1_=0.72) and also for UCYN-A2 (*ρ*_A2_=0.68), as were photosynthesis genes (*ρ*_A1_=0.68, *ρ*_A2_=0.78), and ATP synthases (*ρ*_A1_=0.91, *ρ*_A2_=0.91). Interestingly, the pathway correlations for UCYN-A1 were mainly lower than for UCYN-A2, consistent with the overall Spearman correlations for gene ranks, which might indicate that UCYN-A1 or its host was more impacted by habitat changes than was UCYN-A2 or its host. For example, UCYN-A1 had lower cross-site correlations for N_2_ fixation (*ρ*_A1_=0.54, *ρ*_A2_=0.97), the pentose phosphate pathway (*ρ*_A1_=0.41, *ρ*_A2_=0.64) and RNA degradation genes (*ρ*_A1_=0.11, *ρ*_A2_=0.91). For RNA degradation genes the striking difference was due to *groEL* (GID.646530537 GID.2528847911 in UCYN-A1 and A2, respectively), which were highly transcribed across sites for UCYN-A2 but only at Scripps Pier for UCYN-A1. Most cyanobacteria (including UCYN-A1 and A2) have two *groEL* genes (50, 51), which encode chaperonins within which proteins can refold, alleviating the aggregation of denatured proteins under conditions of heat and other stresses (52). In a culture study of a cyanobacterial diazotroph, overexpression of *groEL* was shown to support increased N_2_ fixation during salt stress (53). Possibly the lower *groEL* transcript levels for UCYN-A1 compared to A2 at Stn. ALOHA are due to different or faster responses to light or temperature at 45 m. Porphyrin and chlorophyll metabolism genes were weakly correlated across sites for both sublineages (*ρ*_A1_=0.30, *ρ*_A2_=0.20). If cross-habitat transcriptomic data were available for the hosts, we could compare host and symbiont pathways that responded to habitat change. This would provide clues as to whether the UCYN-A responses we observed were due to habitat or were host-mediated.

### Transcripts peaked at different times for Stn. ALOHA sublineages, except for genes involved in N_2_ fixation

We looked at the times of day of highest (“peak”) and lowest transcript levels for all detected genes. At Stn. ALOHA peak times differed by sublineage and were mainly at sunset for UCYN-A1 (21% at L12), near sunrise for UCYN-A2 (54% from D9 to L3), and at sunrise or sunset for UCYN-A3 (20% at D12, 19% at L12; Figs 3 and S4). Curiously, all three UCYN-A sublineages had increasing percentages of genes peaking from night through sunrise, but with a 3 h lag between sublineages: Increases started at D3 for UCYN-A2, at D6 for UCYN-A3, and at D9 for UCYN-A1 (Fig S4). In contrast, at Scripps Pier genes tended to peak for both UCYN-A1 and A2 in the late afternoon (L9, 48% for UCYN-A1, 44% for UCYN-A2; Figs 3 and S4). Looking at the 760 cross-site detected genes, peak times changed from Scripps Pier to Stn. ALOHA for most genes from UCYN-A1 (82% of genes) and UCYN-A2 (86%), often by 6 h or more (47% and 51%, respectively). Photosystem I gene *psaA* peaked at midnight (D6) at Scripps Pier but at midday (L6) at Stn. ALOHA for both sublineages. Moreover, the times of lowest transcript levels varied by sublineage at Stn. ALOHA but were mainly at midnight at Scripps Pier (D6, 83% for UCYN-A1, 73% for UCYN-A2; Fig S5). Altogether, these results suggest that conditions of lower light (45 m, 14-19% of surface PAR; (54)) and nutrients at Stn. ALOHA, compared to Scripps Pier, drove UCYN-A sublineages to have different times for highest as well as lowest transcriptional activity for most genes.

**Fig 3.**
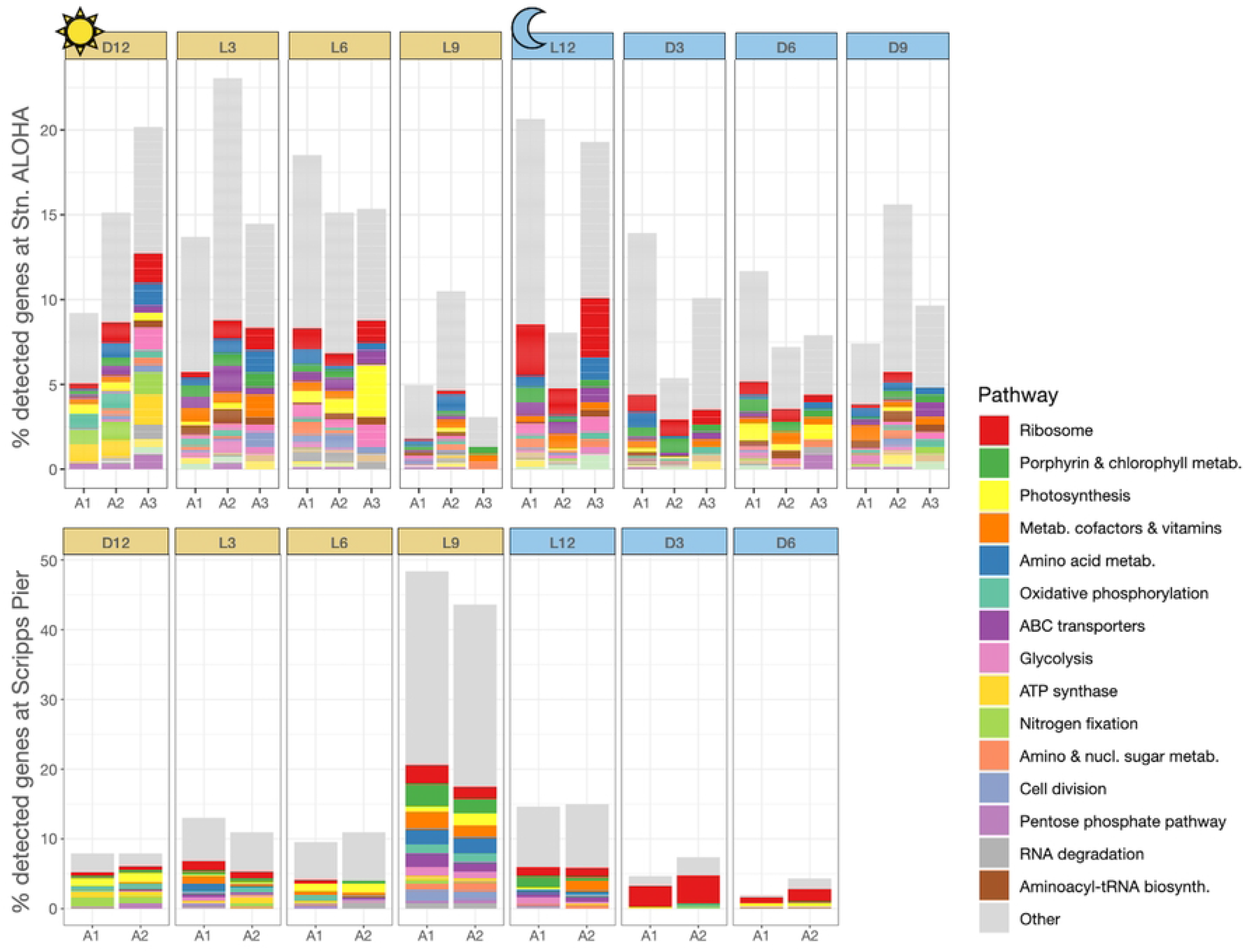
All detected genes at Stn. ALOHA and Scripps Pier are shown at the time of day when their transcript levels peaked. The *x* axes are partitioned first by peak time, then by UCYN-A sublineage. The *y* axes show the proportion of detected genes from the sublineage.

The exceptions were 122 cross-site genes from UCYN-A1 and A2 (*n_A1_*=52, *n_A2_*=70) that had peak transcript levels at the same time of day at Stn. ALOHA and Scripps Pier (S2 Table). For both sublineages many of these genes had roles in N_2_ fixation or photosynthesis, including cytochrome b_6_f complex, ferredoxins, ATP synthases, oxidative phosphorylation, and the metabolism of porphyrin and chlorophyll (but not photosystem I genes). These same pathways were identified above as highly transcribed at both sites. The maintained peak times across sites further support that N_2_ fixation is fundamental across UCYN-A symbioses and environments.

Lower light and nutrients might have contributed to the timing differences, but the exceptions just described suggest that host interactions were important. First, UCYN-A genomes share mostly the same genes among their highly streamlined genomes (7). Moreover, the vast majority of protein coding genes shared by UCYN-A1 and A2 are thought to have evolved under purifying selection within the respective hosts, with niche adaptation (positive selection) attributed to none of the shared genes (46). Second, genes critical to the symbiosis (the exchange of N for C) were the exceptions which maintained the same schedule across sites and UCYN-A sublineages (N_2_ fixation genes as well as ATP synthase genes), which parallels morning peak times for carbon fixation / metabolism genes from haptophytes in the NPSG (55, 56) and California coast (49). Third, at Stn. ALOHA Gradoville et al. (2021) concluded that light did not impact UCYN-A1 nitrogenase transcription because *nifH* transcript levels did not change from the surface to 100m (though N_2_ fixation rates decreased), and they observed morning peaks for UCYN-A1 *nifH* transcript levels across depths (57) as in previous studies (12, 58). Thus, it seems more plausible that the hosts could drive the striking differences in timing for most genes yet maintain morning peaks for genes that underpin the exchange of C and N across different nutrient and light conditions. Host-mediated or not, our results provide evidence for a different regulatory mechanism for genes thought to be essential to UCYN-A symbioses.

### At Stn. ALOHA UCYN-A had fewer diel genes and transcripts peaked more often at sunrise or sunset compared to Scripps Pier

Although many genes were detected at Stn. ALOHA, only ~11% had significantly periodic (“diel”) transcript levels, or 8%, 7%, and 8% of total known genes for UCYN-A1, A2, and A3, respectively (Fig 4, S1 Table). In contrast, at Scripps Pier ~85% of detected genes from UCYN-A1 and A2 were diel (at least 27% of total known genes for each) (16). The substantially lower percentages of diel genes in the present study persisted after controlling for different numbers of time points and detected genes in comparison to the coastal study (Supplemental Information).

**Fig 4.**
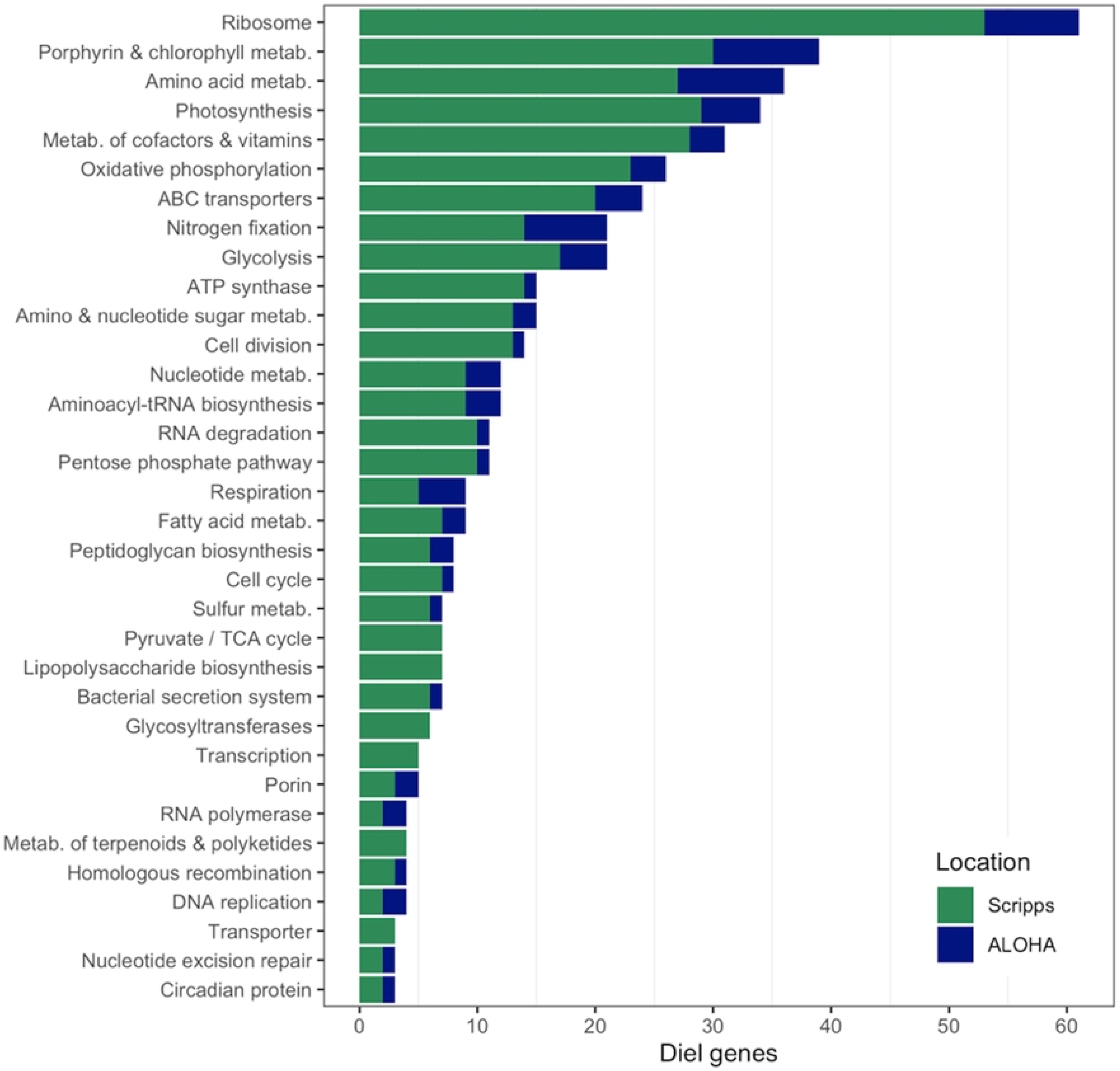
Abundance of UCYN-A1 and A2 diel genes at Stn. ALOHA (blue) and Scripps Pier (green). The *x* axis shows the number of diel genes detected from each pathway. For legibility only pathways with ≥3 diel genes are shown (450 of 459 total diel genes at either site).

Several factors may explain why Stn. ALOHA had far fewer diel genes. The coastal samples were from the surface and likely experienced higher daily integrated light fluxes than Stn. ALOHA samples, which were from 45 m (14-19% of surface PAR; Letelier et al. 2004); total hours of light and dark were similar for both studies despite different seasons. At Stn. ALOHA, Vislova et al. (2019) observed clear decreases in percentages of diel genes from 25 m to 75 m, especially for haptophytes *[under conditions of shallow (<75m) mixing]*. Therefore, they attributed the overall proportions of diel genes *[and the decreases with depth]* to daily integrated light flux. Other factors may have contributed to the differences at Stn. ALOHA and Scripps Pier. Waters near Hawaii experience high levels of sunlight and warm temperatures year round while coastal California waters are colder and undergo marked seasonal transitions. Nutrient composition and organismal interactions may also be factors: Light may drive the release of organic matter by photosynthetic organisms at Stn. ALOHA, but land-derived organic matter is an additional factor at Scripps Pier.

Next we checked whether the time of peak transcript levels changed for genes that were diel in both the NPSG and the coastal site (“cross-site diel genes”). Although nearly every diel gene at Scripps Pier was detected at Stn. ALOHA (649 of 651 diel genes), only 62 were cross-site diel (*n_A1_*=32, *n_A2_*=30; S3 Table). For both UCYN-A1 and A2, cross-site diel genes that peaked at sunrise (D12) at Scripps Pier also peaked at sunrise at Stn. ALOHA (Fig S6). For UCYN-A1 these included genes for nitrogen fixation (*nifBESU*), the circadian oscillating protein 23 (COP23), and the metabolism of porphyrin and chlorophyll, which also always peaked at sunrise for UCYN-A2 (Fig S6). Sunset (L12) peaks also usually persisted from Scripps Pier to Stn. ALOHA (for 5 of 6 UCYN-A1 genes (Fig S6); UCYN-A2 had no cross-site diel genes that peaked at sunset; S3 Table). These included genes for the metabolism of amino acids (*glnA*), amino sugars/nucleotides (SQD1), and vitamin E. Interestingly, for both sublineages cross-site diel genes most often peaked in the afternoon (L9) at Scripps Pier (17 genes for UCYN-A1, 12 genes for UCYN-A2) but rarely peaked in the afternoon at Stn. ALOHA (one UCYN-A1 gene predicted to regulate disulfide bond formation) (S3 Table, Figs 3 and S4). For UCYN-A1, peaks often shifted from the afternoon (L9) at Scripps Pier to sunset at Stn. ALOHA, including genes for porphyrin and chlorophyll metabolism, nucleotide metabolism (*add*), and a ribosomal protein (RP-S4). A remarkable exception was the ATP synthase gene ATPF0C whose peak shifted from the afternoon at Scripps Pier to sunrise (D12) at Stn. ALOHA, synchronous with *nif* gene peaks. For UCYN-A2, 4 of the cross-site diel genes had peaks shift from the afternoon at Scripps Pier to sunrise at Stn. ALOHA, including genes for glycolysis (FBA) and the metabolism of cofactors and vitamins (*thiL*). Six of the remaining 8 genes had peaks shift to the morning (RNA degradation gene (*rnr*)) or noon (peptidoglycan biosynthesis [*vanY*], ferredoxin [*petB*]). Altogether these results suggest that for both UCYN-A1 and A2 the different conditions (e.g. light and nutrients) at Scripps Pier and Stn. ALOHA led to fewer genes with significantly periodic expression, and to shifts from mainly afternoon peaks at the coast to sunrise or sunset peaks at Stn. ALOHA.

These different conditions could affect UCYN-A directly or through the host. For example, glycolysis genes for UCYN-A1 showed diel patterns at the Scripps Pier but not at Stn. ALOHA (Fig 4). Possibly UCYN-A1 glycolysis genes tracked carbohydrates supplied by the host, which may have been from photosynthesis (periodic) at Scripps Pier but heterotrophy (aperiodic) at Stn. ALOHA at 45 m.

## Conclusions

This study provides the first comparison of metatranscriptomes from natural populations of UCYN-A in oligotrophic and coastal waters. Across environments and sublineages, UCYN-A had the same highly transcribed genes from indispensable pathways including photosynthesis, ATP synthases, N_2_ fixation, and respiration. Moreover, under conditions of lower light and nutrients UCYN-A maintained morning peaks for genes that underpin the exchange of fixed N_2_ with C from the host.

On the other hand, UCYN-A sensitivity to habitat was illustrated by the striking differences in transcription at coastal and oligotrophic sites. At the coastal site transcription patterns were similar across sublineages, but at Stn. ALOHA most genes had peaks shift, often to near sunrise or sunset, and sublineages took on distinct daily transcription schedules. The complex sublineage-specific responses at Stn. ALOHA, but not at the coastal site, point to host interactions as important drivers of UCYN-A transcriptomes. Further studies are needed to understand the regulation and detailed functions of genes that were differentially expressed between sublineages and sites. We hope that the patterns and responsive genes identified in the present work will be targets for studies of UCYN-A symbioses in culture, which has only recently become possible. Culture studies could show whether environmental changes impact host circadian rhythms, including which genes and metabolites might be affect the daily transcription cycle in UCYN-A.

## Acknowledgments

We are grateful to the captain and crew of the R/V Kilo Moana for their essential support during C-MORE Cruise C-20. M.C.M.-M. designed the experiment and the UCYN-A microarray and collected and prepared all samples. J.D.M analyzed the data and J.M prepared the figures. M.C.M.-M., J.D.M., and J.P.Z. drafted the manuscript. All authors read and approved the final manuscript.

## Supporting information captions

**S1 Fig. BLAST-based simulation of hybridization to probes.**

*In silico* hybridizations simulated Agilent SurePrint technology. Hybridization occurred if the BLAST high-scoring pair (HSP) aligned at >95%id and >95% of the 60 nt probe length.

**S2 Fig. Distributions of raw microarray probe intensities for each UCYN-A sublineage.** Includes only probes that are <95%id to any other probe (the vast majority) and therefore not expected to cross-hybridize among sublineages detected by Agilent SurePrint microarrays. The UCYN-A2 distribution suggests that UCYN-A2 transcripts were highly abundant, despite it being only ~0.7% of the UCYN-A population at Stn. ALOHA. *n* indicates the numbers of probes for each sublineage.

**S3 Fig. Correlations across sites of two UCYN-A sublineages.**

For each UCYN-A sublineage and pathway, the Pearson correlations across sites (NPSG and coastal) were calculated for the median transcript levels for the genes in the pathway (requiring ≥5 genes detected at both sites). The plot shows that 7 pathways (discs) had significantly correlated transcript levels (*p*<0.05) across sites for both UCYN-A1 and A2. Another 5 pathways (triangles) were significantly correlated across sites only for UCYN-A2.

**S4 Fig. At Stn. ALOHA, UCYN-A sublineages had different schedules for peak transcriptional activity.**

As in Fig. 3, all detected genes were categorized by the time of their peak transcript level. At Stn. ALOHA, UCYN-A sublineages had a 3 h lag between the times when many of their genes peaked: D12 for UCYN-A3, L3 for UCYN-A2, and L6 for UCYN-A1. No sample was collected at D9 in the Scripps Pier study so that time point lacks an open circle.

**S5 Fig. Times of day at which all detected genes from UCYN-A sublineages reached their lowest transcript levels at Stn. ALOHA and at Scripps Pier**.

The *x* axes are partitioned first by peak time, then by UCYN-A sublineage. The *y* axes show the proportion of detected genes from the sublineage.

**S6 Fig. Diel genes at Stn. ALOHA mainly peaked at sunrise (D12) or sunset (L12), unlike at Scripps Pier.**

At Stn. ALOHA clustering assigned 154 of the 188 diel genes to the sunrise peak cluster (68 genes) or the sunset peak cluster (86 genes). For legibility only 53 diel genes are shown. Genes are colored by their diel cluster in the study at Scripps Pier (16). Within each of the four plots genes with different colors appear, which indicates that genes with different diel schedules (clusters) at Scripps Pier changed to have the same diel schedule at Stn. ALOHA. At Scripps Pier 389 of the 651 diel genes were in cluster I (199 genes, L9 peak) or cluster II (190 genes, D12/L3 peak).

**S1 Table. Detected genes at Stn. ALOHA.**

For each of the 1939 detected genes at Stn. ALOHA, the normalized log_2_ transcript levels, annotation, and Fourier analysis results are provided.

**S2 Table. Genes detected at both Stn. ALOHA and Scripps Pier.**

The 122 cross-site detected genes from UCYN-A1 and UCYN-A2 are listed with annotation.

**S3 Table. Comparison of genes that had significant periodic (“diel”) expression at both Stn. ALOHA and Scripps Pier.**

The first sheet categorizes the cross-site diel genes by the time of day at which they peaked at Stn. ALOHA versus at Scripps Pier. The second and third sheets describe the cross-site diel genes from UCYN-A1 and UCYN-A2, respectively.

